# Distinctive desmoplastic 3D morphology associated with BRAF^V600E^ in papillary thyroid cancers

**DOI:** 10.1101/160127

**Authors:** M. Tarabichi, A. Antoniou, S. Le Pennec, D. Gacquer, N. de Saint Aubain, L. Craciun, T. Cielen, I. Laios, D. Larsimont, G. Andry, J.E. Dumont, C. Maenhaut, V. Detours

## Abstract

Textbooks suggest that cancers are compact balls with an inner core and an invasive front in contact with non-cancerous cells, and that tumor growth is driven by tumor cell proliferation subsequent to oncogenic genome mutations. We reconstructed at histological scale the 3D volume occupied by tumor cells in two regions of a BRAF^V600E^-mutated papillary thyroid carcinoma (PTC) with low tumor purity, as determined by sequencing, but initially considered high purity during pathology review. In contrast with a compact ball, tumor cells formed a sparse mesh deeply embedded within the stroma. The concepts of inner core and invasive front broke down in this morphology: all tumor cells were within short distance from the stroma. The fibrous stroma was highly cellular and proliferative, a result confirmed in an independent series. This case was not unique: 3.5% of The Cancer Genome Atlas PTCs had purities <25%, 27% were below the 60% purity inclusion criterion. Moreover, the presence of BRAF^V600E^ was associated with extensive fibrosis, high stromal activation, and dedifferentiation. Thus, non-tumor cells make most of the tumor mass and contribute to its expansion in a significant fraction of BRAF^V600E^ PTCs. Therapeutics targeting the tumor-stroma crosstalk could be beneficial in this context.

## Introduction

In order to focus on informative sequencing data most oncogenomic studies reject samples with low tumor cell content as estimated beforehand from tissue examination by board-certified pathologists. Nevertheless, low tumor purity samples routinely end up being sequenced, and in most cases, are deemed useless for further analysis. That tumor purity is not necessarily reliably estimated from the expert observation of tumor morphology is intriguing. This paper stems from the unwanted sequencing of such series of tumor regions, which were originally collected for a study of the intra-tumor and inter-foci heterogeneity in a papillary thyroid cancer (PTC) patient. Low purity was obviously incompatible with our original goal. But instead of shelving the data, we reasoned that further investigation of the underlying tumor morphology could shed light on non-cancer cells contribution to tumor expansion.

The shape of the tumor mass is important as it defines the contact area with the stroma, where features of aggressiveness such as invasiveness and increased proliferation are observed [1], some of which carry prognostic information in various cancer types, e.g. in oral cancers [2]. In fact, together with the morphology of the tumor cells, the morphology of the tumor mass relates to the mode of metastatic dissemination [3]. Notably, cancer-associated fibroblasts (CAF) have been shown to pull tumor cells, generating tumor elongations [4]. Interestingly, many PTC present poorly defined boundaries [5]. Liu *et al.* showed an association between PTC recurrence and presence of isolated clusters of cells and the loss of cell polarity, i.e. two features of EMT [6]. Moreover, microPTC^V600E^ (PTC<1cm, with BRAF^V600E^ mutation) seemed to present more complex architectural invasive fronts, with markedly invasive contours than microPTC^WT^ [7]. Isolated clusters of tumor cells at the invasive front could be interpreted as the consequence of an EMT followed by a return to the epithelial state [1]. Alternatively, these isolated islands of cells observed on 2D slices could also belong to the same connected 3D tumor volume, implying branching patterns drifting from an ellipsoid shape.

So far, 3D tumor volumes have been mostly acquired by medical imaging techniques, such as computed tomography and magnetic resonance imaging, common in the field of radiomics [8]. These technologies, which operate at macroscopic scale, typically yield ellipsoid-shaped tumor volumes with fingering patterns. In this context the tumor/stroma interface is implicitly assumed to be located on an ellipsoid periphery. For example, Vasko and collaborators analysed differential expression between peripheral and central parts of PTC, discussing a higher epithelial-to-mesenchymal transition (EMT) at the periphery [9]. The assumption that the tumor-stroma contact surface is ellipsoid, however, is far from granted if one consider the fine-scale relevant to cell-to-cell contact.

When zooming in the invasive front with higher resolution imaging, the observed tumor-stroma contact area might increase quite a lot, akin to coastline lengths at different magnifications in the coastline paradox [10]. The fractal dimension measures the increase in tumor-stroma contact area in proportion to the magnification. At the higher histological resolution, complex, non ellipsoid, morphology of the tumor mass have been described and their fractal dimensions were linked with prognosis [11]. Tumor budding in colorectal carcinoma is another complex pattern whereby isolated clusters of cells at the invasive front would have gone through EMT, sign of a worse prognosis [12]. In our study, we explore the shape of the tumor/stroma contact area at histological scale using advanced 3D imaging.

Papillary Thyroid Carcinomas (PTC) represent ∼80% of all thyroid cancers [13] and, given the fast increasing incidence of microPTC, which can be ascribed mostly to earlier diagnosis, thyroid cancer, could become the fourth most diagnosed cancer in the United States by 2030, ranked before colorectal cancer [14].

A recent study by *The Cancer Genome Atlas* (TCGA) [15] could identify potential driver mutations for 96.5% of 496 PTC. In line with previous studies PTC progression was essentially driven by constitutive activation of the MAPK pathway, with the most frequent mutations found in the *BRAF, RAS* and *RET* genes [15], [16]. BRAF^V600E^ was by far the most frequent mutation, affecting ∼60% of PTC [15], [17]. This mutation is also found in many other cancer types, among which melanomas and colorectal cancers.

Although many studies showed association of BRAF^V600E^ with clinical features of aggressiveness and with bad prognosis in PTC (e.g. [18] [7]), this has been often disputed [19]. Most notably, BRAF^V600E^ was not associated with distant metastasis [20], [21]. However, in PTC^V600E^ presenting poorly-differentiated or undifferentiated areas it was consistently detected in both regions, and might thus play a role in the progression from PTC to the aggressive undifferentiated cancers [17], [22]. Here, the joint analysis of TCGA histological imaging, genomic and transcriptomic data reveals that our low purity multi-region PTC is not unique. It is part of continuum of PTC morphologies in which BRAF^V600E^ PTC stand out with strong CAF activation, lower tumor cell purity, dedifferentiation of tumor cells, and high proliferation rate in both the tumor cell and the CAF compartments.

## Results

*Multi-region sequencing and imaging reveal a low cellularity in ∼30% of BRAF^V600E^ papillary thyroid cancers, which escapes pathologists’ examination.* We performed multi-region exome sequencing of a BRAF^V600E^ PTC (sampling protocol Figure 1A-B; case description in Material and Methods). The regions included five primary tumor blocks, 4 nodal metastases, and as controls, three histologically normal thyroid blocks and a normal lymph node.

**Figure 1.**
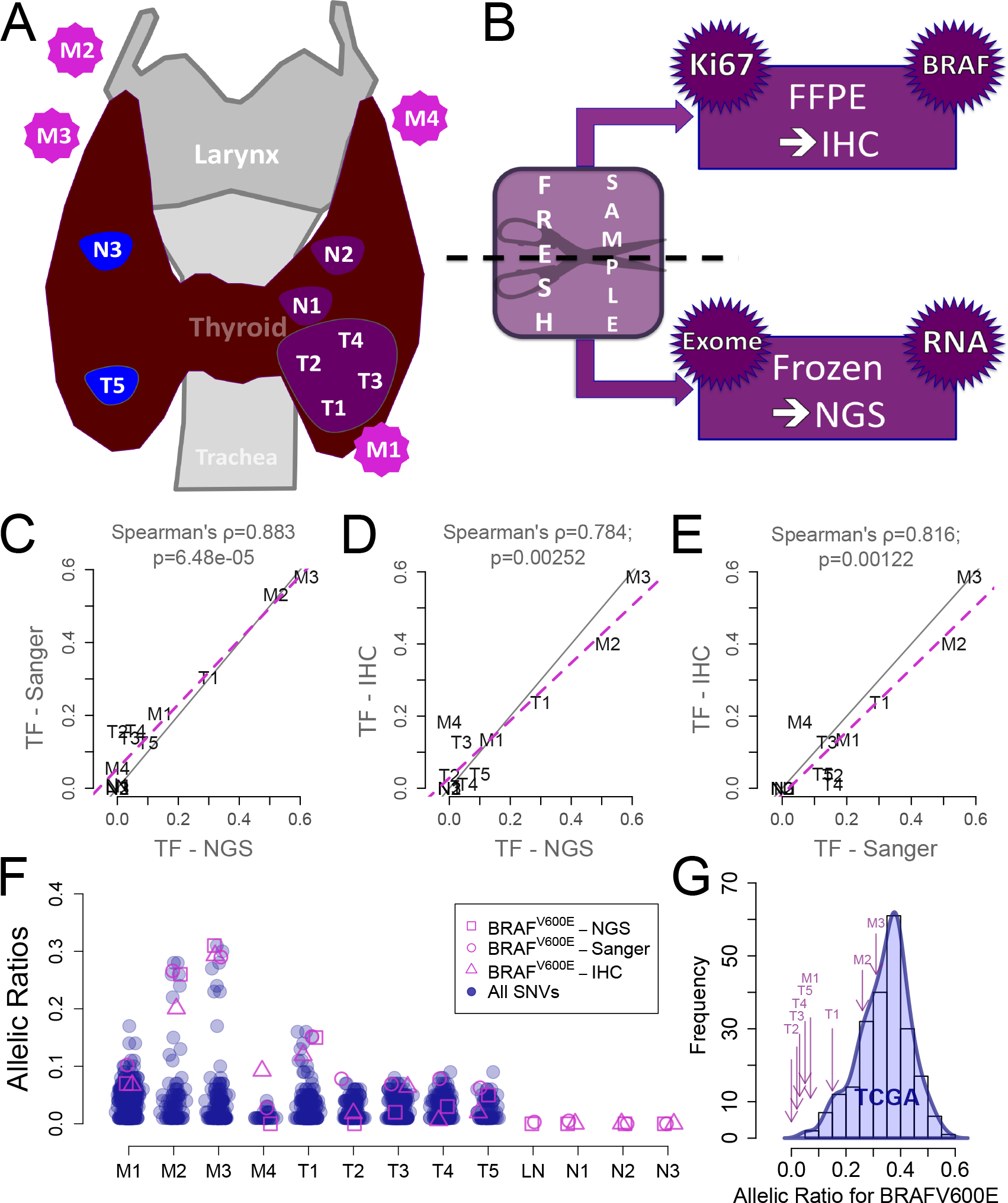
Heterogeneity in tumor purity, BRAF^V600E^ clonality and protein expression in a PTC patient. A. Samples localization in the patient (T for primary tumor block; N for normal adjacent thyroid tissue; M for nodal metastasis). B. Material preparation: each block was halved, one part was frozen for exome- and RNA-sequencing (not used in this study), the other embedded in paraffin (FFPE) for immunohistochemistry (IHC) staining for BRAF^V600E^ and Ki67. C. Tissue fractions (TF) quantifications (twice the allelic ratios) of BRAF^V600E^ in the DNA of our blocks: Sanger *vs.* next-generation sequencing (NGS). D. For each block, BRAF^V600E^ NGS TF *vs.* protein expression ratio computed from IHC images. E. Same as E for Sanger TF against protein expression ratios. F. For each block, allelic ratios of all somatic mutations (blue) against the three independent measurements of BRAF^V600E^ ratios (in pink: Sanger, NGS and IHC ratios). G. Distribution of allelic ratios for BRAF^V600E^ in TCGA PTC (blue) and allelic ratios for our patient’s blocks (purple arrows).

The allelic ratio of the BRAF^V600E^ mutation estimated from sequencing data ranged from 0 to 30%, corresponding to 0 to 60% tissue fractions (Figure 1C). Quantitative analysis of Sanger sequencing technically validated these allelic ratios (Figure 1C). Such low ratios could have resulted either from low tumor cell fractions, copy number aberration or from subclonal BRAF^V600E^ mutations.

Subclonality of the mutation was suggested in previous studies [23], [24], To address this possibility, we reasoned that subclonal BRAF^V600E^ mutations are incompatible with an initiation of tumor growth by BRAF^V600E^. Thus, if BRAF^V600E^ mutation were subclonal, then other mutations must have initiated the tumor, i.e. be the ancestor of all the tumor cells, and have a greater allelic fraction than BRAF^V600E^ in all the regions. To test this prediction, we compared in each region the allelic ratios of BRAF^V600E^ and of all other mutations (Figure 1F). In all regions but one, BRAF^V600E^ was among the mutations with the largest allelic ratios. This ruled out the subclonality of BRAF^V600E^ in our samples.

Having established clonality, we assessed the alternative hypothesis that the low BRAF^V600E^ allelic ratios reflected a low fraction of tumor cells within the tumor mass. Paraffin-embedded slices immediately adjacent to each one of the tumor regions used for exome sequencing were stained with an antibody specific of the V600E-mutated BRAF protein (Figure 1AB; Figure 2A). Taking advantage of the distinctive hue of stained cells we devised an image-processing pipeline (Material and Methods) that estimates automatically and quantitatively the areas occupied by BRAF^V600E^ tumor cells and by the entire tumor slice. We then compared the tumor cell area ratio to the tissue fraction estimated from their region-matched BRAF^V600E^ allelic ratio (Figure 1D and E). They were highly correlated, strongly supporting that the low allelic ratios observed in sequence data resulted from a low ratio of tumor cells within the overall tumor mass.

**Figure 2.**
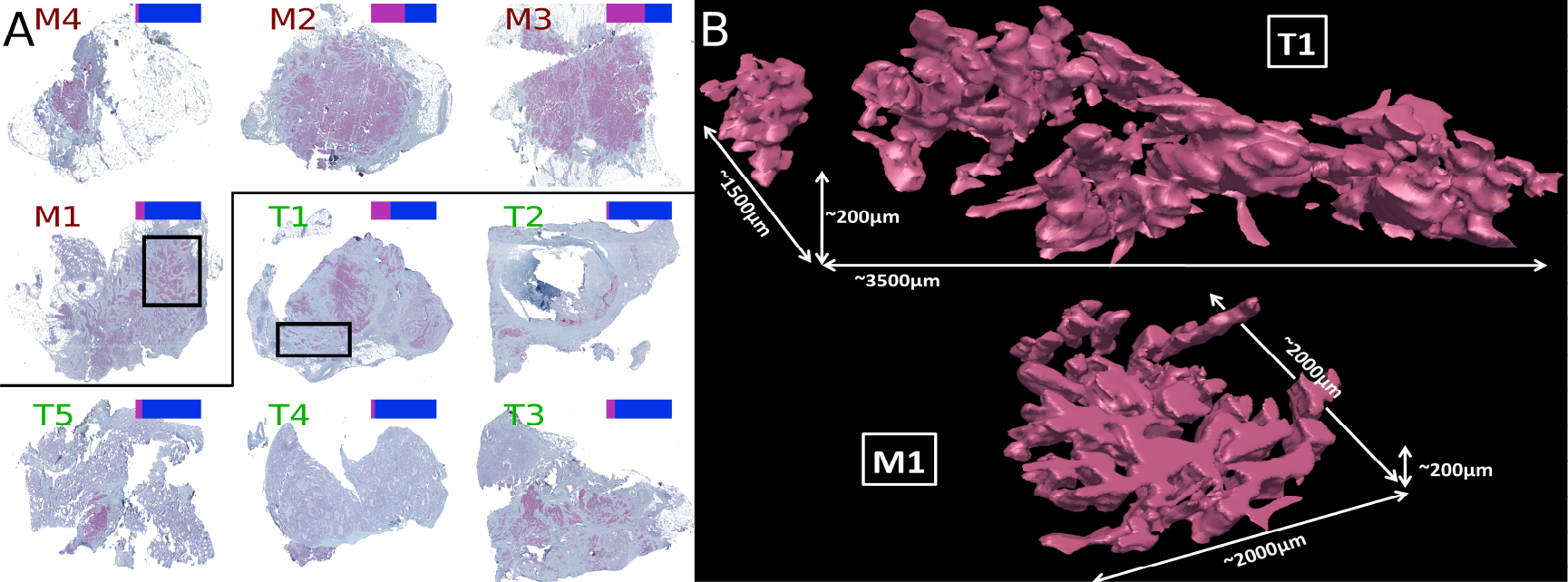
Tumor shape of a PTC in 2D and 3D at histologic resolution. A. For each tumor sample a 2D slice stained for BRAF^V600E^ is shown. M1-4 are the nodal metastases; M2-4 are recurrences. T1-5 are the primary tumors. On the top right of each image a color bar shows the proportion of marked (pink) vs. unmarked (blue) regions as extracted by image analyses. B. 3D reconstruction for the T1 primary tumor and the M1 nodal metastasis. Dimensions are given for each part. For T1 one isolated island on the left was found, whereas in M1, all cells belong to the same connected component. In both blocks the tumor part forms a complex 3D network infiltrating the stroma. Supplementary videos S1 and S2 show rotations of the 3D model of T1.

This result was surprising given that the samples had been sent beforehand for a pathology review to guarantee a cellularity greater than 70%. Indeed, we initially generated the sequencing data for a genomic intra-tumor heterogeneity study, which required a minimum tumor purity to control the power to detect subclonal variants at a given coverage. But cellularity had obviously been grossly underestimated. As it turned out, the vast fibrotic regions present in the tumor mass (Figure 2A) were unexpectedly cellular.

How commonly is cellularity underestimated? To address this question we obtained allelic ratios for the mutation corresponding to BRAF^V600E^, as measured from exome sequencing data from the 211 PTCs of the TCGA [15] (Figure 1G). As expected, our tumor regions were in the lower end of the TCGA purity spectrum, but not below it. Moreover, the *a priori* pathologic purity threshold for the inclusion of PTC samples in the TCGA was 60%, whereas the purities computed *in silico a posteriori* were <60% for around one third of the samples (Supplementary Figure S1). We concluded that tumor cell purity is low in a nontrivial fraction of PTCs and that it is often underestimated during pathology examination.

*3D histological scale reconstruction of low purity BRAF^V600E^ papillary thyroid cancer blocks reveal dense connected fractal-like meshes of tumor cells, with an extensive contact surface with the fibrous stroma.* We asked why low purity PTCs were much more frequent than estimated during pathology review, and conducted an in-depth analysis of our PTC samples. In all regions, BRAF^V600E^-staining revealed isolated patches of tumor cells embedded within an extensive stroma (Figure 2A). Cells that appear to be isolated in two dimensions, however, could turn out to be part of the same tumor mass when observed in three dimensions.

A novel protocol was devised to reconstruct and visualize the geometry of tumor cell aggregates in 3D (Figure 2A; material and methods). First, tumor blocks were serially sliced. Second, each slice was stained for BRAF^V600E^. Third, the contours of stained areas were determined in each slice images. Fourth, images and contours were registered manually and locally within the small volume of interest in order to minimize discrepancies arising from tissue deformation. Fifth, the 3D envelops of the tumor cell were computed from the stacked 2D contours.

This 3D reconstruction at histological scale was applied to a nodal metastasis block (M1, Figure 2AB, Supplementary Video S1) and a primary tumor block (T1, Figure 2AB, Supplementary Video S2) of about 1mm^3^ each. For block M1, no isolated island of tumor cells was found in the tumor volume reconstructed over a depth of 200μm, i.e. all cells were part of a single connected component (Figure 2B). For block T1, the number of disconnected tumor cell clusters was reduced from a dozen in the 2D slice to 2 in the 200μm-deep 3D microvolume (Figure 2B). This strongly suggested that the tumor cells within a region formed a single connected component.

Interestingly, the 3D volumes departed drastically from any ellipsoidal shape. Tumor cells formed a complex volume deeply embedded within the stroma (Figure 2B). Instead of a tumor with an invasive front in contact with the stroma and inner tumor cells in contact mostly with other tumor cells, we observed that a the vast majority of tumor cells were in direct contact with, or nearby, the stroma (Figure 2B).

*Stromal cells and cancer cells proliferate at comparable rates in PTCs.* The results of previous sections raised a fundamental question: from the mixture of stromal and tumor cells, which cell types drive tumor growth?

We performed Ki67 staining on our regions and compared proliferation rates of normal stroma, stroma adjacent to tumor, non-cancerous follicular thyroid cells and tumor cells (Figure 3A). As expected, proliferation rates were higher in tumor blocks than normal thyroid blocks. Primary tumor regions and metastases had similar proliferation rates. Importantly, the proliferation rate of stromal fibroblasts was equal to, or higher than, that of follicular thyrocytes. Statistical significance, however, could not be reliably established with this limited dataset.

**Figure 3.**
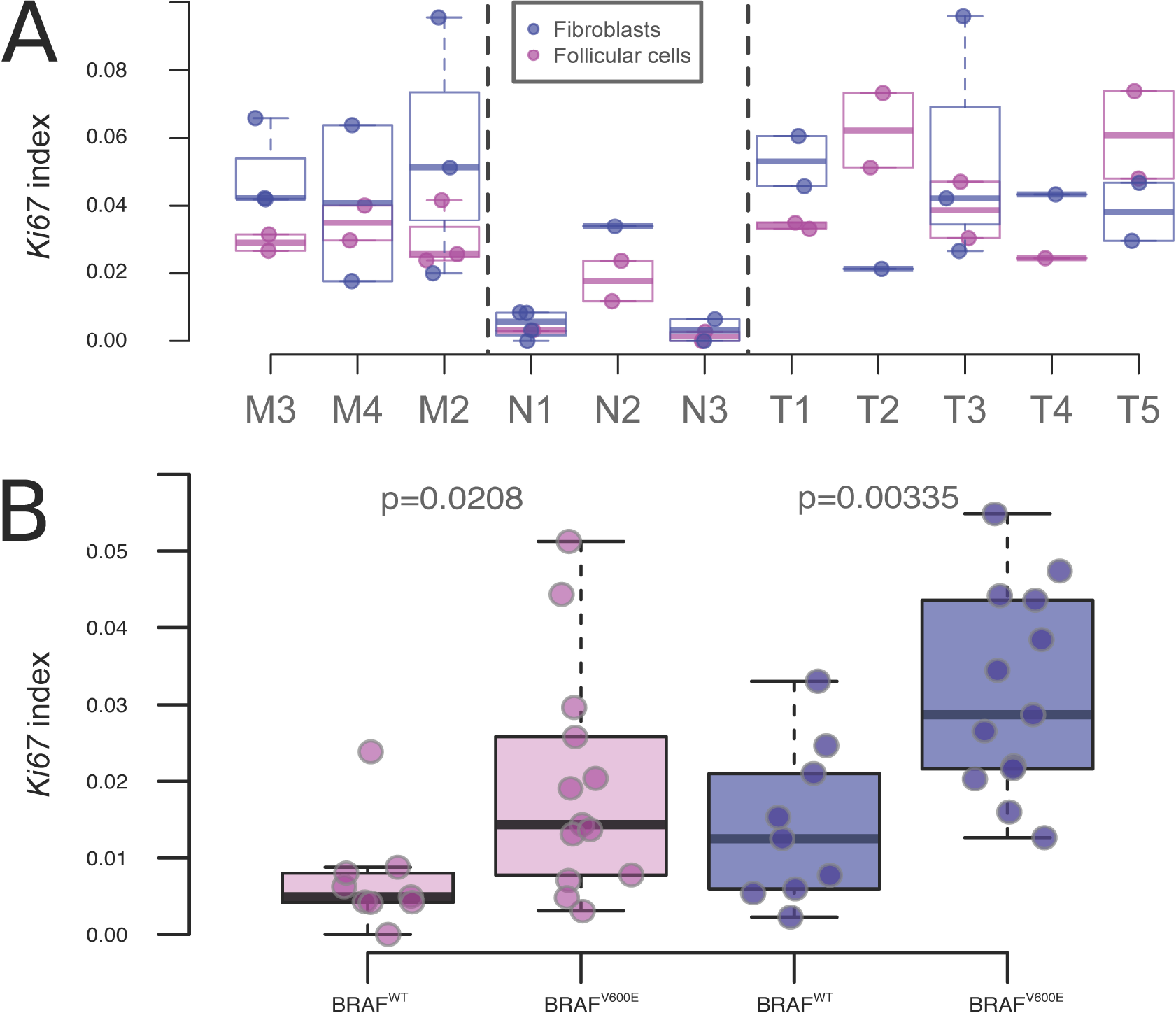
Proliferation of epithelial cells and fibroblasts. A. For each block, Ki67 proliferation index was derived for cells of thyroid and stromal background in at least one part of the block. The Ki67 ratios are shown on this boxplot for each investigated block in stromal (blue) and thyroid (pink) parts. B. Ki67 indices for an independent series of 22 PTCs, of which 13 were BRAF-mutated. Each point is an average of two Ki67 values for two independent regions on the slices.

To generalise this finding, we measured Ki67 indices in the stromal and tumor compartments of 22 independent PTCs, including 13 BRAF^V600E^ and 9 BRAF^WT^ tumors (Figure 3B). Importantly, these samples were not selected for specific tumor cell purity. For each tumor, one slice was stained for Ki67. The ratio of Ki67-stained nuclei for tumor cells was not higher than for stromal cells, suggesting that the density of cycling cells in the fibrous stroma was high.

*BRAF^V600E^ PTCs are associated with a desmoplastic phenotype, a higher proliferation rate and a decreased differentiation.* Interestingly, Ki67 staining also revealed a higher fraction of tumor and stromal cycling cells in BRAF^V600E^ than in BRAF^WT^ PTCs (Figure 3B). We combined image, genomic and transcriptomic analyses of the 496 PTCs from the TCGA [15] to further characterize the phenotype of BRAF^V600E^-mutated PTC and how it relates to the relative sizes of the tumor and stromal compartments.

We have shown in Figure 1 how to estimate the tumor cell fraction from histological slices using image analysis. The estimates were confirmed by comparisons with the BRAF^V600E^ allelic fraction, thereby validating the method. To extend this analysis to the TCGA cohort, we downloaded the PTC histological H&E slide images from the TCGA cancer digital archives (http://cancer.digitalslidearchive.net). Because fibrosis presented a distinctive colour hue (Figure 4A), it was possible to automatically estimate fibrotic content as the ratio of the fibrotic areas to the total tumor area. Fibrotic content was higher in PTC^V600E^ than in other PTCs (Figure 4B). RAS mutated tumors had the least fibrous tissue.

**Figure 4.**
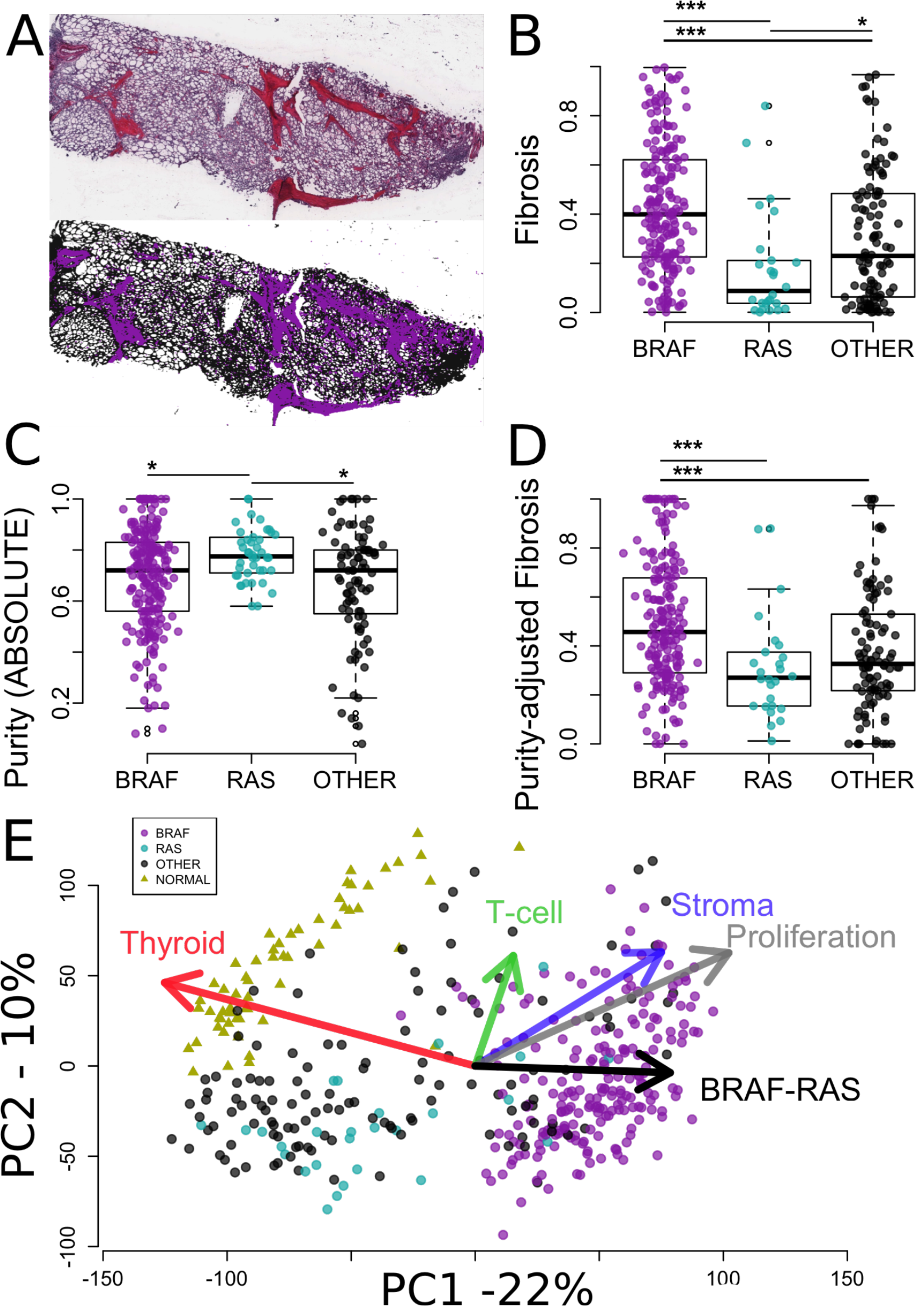
PTC^V600E^ gene expression, purity and fibrosis phenotypes in TCGA PTC. A. At the top, histological image of one PTC from TCGA, stained used H&E (http://cancer.digitalslidearchive.net/), where fibrosis appears with a pink color hue distinct from the blue color hue of surrounding tissues. At the bottom, selection of fibrosis (purple) and surrounding tissues (black) based on their respective hues. The fibrosis content was calculated from these two areas. B. Fibrosis content by mutational group for all TCGA PTC with available histological images (N=331). C. Purity by mutational group of TCGA PTC as computed by ABSOLUTE in the original publication (N=364) [15]. E. Fibrosis by mutational group after correction for purity of the samples using a robust linear regression model (N=331). D. Projection of 423 transcriptomics profiles and gene signatures (arrows) are projected onto the 2 first components computed from the TCGA PTC (circles) and normal adjacent tissues (triangles). Gene expression signatures are the median values of genes related to thyroid, T-cell and stroma, proliferation, and BRAF-RAS genes (Material and Methods; Supplementary material). Arrows point to the directions of higher expression of the genes present in the underlying signatures.

The area of the fibrotic regions does not necessarily reflect the number of cells within these regions. We therefore investigated tumor purity, as measured from genomic data by ABSOLUTE [25] in the original TCGA publication [15]. Purity was lower in BRAF^V600E^ PTCs than in BRAF^WT^ PTCs (Figure 4C). RAS-mutated PTC had the highest purity among all groups. Thus, the distribution of purity across the three groups mirrored that of fibrosis, i.e. a lower fraction of tumor cells was associated with more fibrosis, supporting the qualitative visual assessment that the fibrotic regions contained many fibroblasts. To further define the activity of these fibrotic regions we corrected the fibrosis ratios for the purity of the samples using a robust linear model (Figure 4D). The association of BRAF^V600E^ with fibrosis remained highly significant. Taken together with the Ki67 data from our cohort, these results suggest that BRAF^V600E^ PTCs are associated with a desmoplastic phenotype: they include more fibroblasts, which proliferate more and produce more fibrosis.

To gain a broader view of the phenotype of individual PTC from the TCGA we derived five gene expression signatures measuring different aspects of PTC biology. Two were derived following the procedure of the original TCGA study [15]. They included genes associated with thyroid differentiation and with the BRAF-RAS axis [15]. We also derived three additional signatures for stroma, T-cells, and proliferation (see Material and Methods). Associations between BRAF mutational status and each one of the five gene expression signatures were statistically significant (Mann Whitney tests; false discovery rates <10^-5^). To visualize the global trends in gene expression, we performed a principal component analysis of TCGA RNA-seq data. Expression profiles were projected on the two first components, together with the five signature vectors (Figure 4F). On Figure 4F, points and triangles depict transcriptomes of PTCs and normal thyroid tissues, respectively. Their relative projection on each signature vector is closely related to the relative average expression of the genes making up the underlying gene signatures. BRAF^V600E^ PTCs grouped separately from normal tissues along axes of thyroid differentiation, proliferation and BRAF-RAS expression differences. As expected from the TCGA study, they further separated from BRAF^WT^ PTCs along the vector representing thyroid differentiation and BRAF-RAS expression differences, but also along the vectors standing for stroma, lymphocytes, and proliferation (Figure 4F). Interestingly, the stroma and proliferation axes were strongly associated in these PTCs, which was reflected in their almost identical directions on Figure 4F. Together the higher proliferation and dedifferentiation indices of BRAF^V600E^ PTCs point to a more aggressive phenotype.

## Discussion

Textbooks and low resolution imaging technologies suggest that cancers expand as a compact near-ellipsoid ball with an inner core and an invasive front in contact with non-cancerous tissues. The histological scale 3D reconstruction of our low purity PTC demonstrated the existence of a morphology departing radically from this archetype. Far from an ellipsoid ball, this morphology would be better depicted as a branching fractal-like aggregate of cancer cells deeply embedded within the stroma. In this morphology, the concepts of inner core and invasive front break down because all tumor cells are within short distance from the stroma. The 3D reconstructions also suggested that tumor cells belonged to a single connected component within the ∼1mm^3^ volumes investigated. Thus, all cancer cells were in contact with or close proximity from both stromal and cancer cells.

CAFs have been shown to lead collective invasion of carcinoma cells [4]. The same phenomenon could apply in PTC, as elongating patterns of neoplastic cells into non-neoplastic cells have been described in microPTC [7]. TGF-β signalling was involved in many tumor-stromal interactions involving EMT, proliferation and activating signals. PTC cells could send signals to its surrounding stroma, potentially involving TGF-β1 [26], with increased response-to-signal ratio in PTC^V600E^.

Our analysis also demonstrates the textbook view that tumor expansion is driven by tumor cells proliferation subsequently to oncogenic mutations in their genomes is not universal. Tumor cells represented a small fraction of the total number of cells making the tumor mass, less than 10% in several regions. How, in such circumstances, can the tumor ever reach a clinically detectable volume? Is excess proliferation an exclusive property of the tumor cells? We measured proliferation rates in the tumor and stomal compartments of our original low purity regions and associated normal thyroid tissues and in an independent series of PTCs and normal thyroid tissues. Importantly, the later were not selected for specific purity or fibrotic content. Proliferation was higher in tumor than in normal tissue for both compartments. Our estimations based on Ki67 staining suggested that fibroblasts proliferate faster than tumor cells (Figure 3B). However, unstained nuclei are more easily detected in cancer cells than in fibroblasts, which may bias any direct comparison of proliferation between these compartments. At this point, we can only assert that the proliferation of fibroblasts is increased in PTC compared to normal thyroid. Further research is needed to quantify precisely the relative contribution of both compartments to tumor expansion.

Pathology examination initially suggested that our samples had purities greater than 70%. Subsequent sequencing and computational analyses demonstrated that they were much lower. Fibrotic regions, it turned out, were more cellular than expected. Similarly, inclusion of samples in TCGA thyroid cancer dataset required a purity greater than 60% upon pathology review, but subsequent sequencing and analysis demonstrated that this criteria was not met by one third of the samples. Purity could be underestimated in other studies. Our samples were at the lower end of the purity spectrum, but they were not unique: 3.5% of the 342 TCGA samples analysed had purities below 25%. Given that the inclusion criteria of the TCGA introduced a selection bias towards higher purity, 3.5% would be a lower bound estimate. Incidentally, the association of low purity with BRAF^V600E^ implies that BRAF^V600E^ incidence could also be underestimated in TCGA and possibly other studies.

To better interpret the genome-derived purity data, we also analysed the fibrotic content of TCGA PTC from tumor images, and the expression of stroma-associated genes. We observed a wide spectrum of purity, fibrosis, and stromal gene expression scores. Within this spectrum, nearly all RAS-mutated PTC had high purity, low fibrosis and low stromal gene expression. By contrast, other tumors, most of them harboring BRAF^V600E^, covered the entire spectrum of scores, including a sizable fraction of tumors with low purity, and high fibrosis and stromal gene expression. Previous literature pointed to a role of CAFs in this lower purity. In one study, PTC^V600E^ showed more with stromal fibrosis/sclerosis/desmoplasia and infiltrative growth [27]. In another study, microPTC^V600E^ also had more fibrosis content [7]. Interestingly, both the image-derived fibrotic and the stromal gene expression scores were associated with purity independently of purity in our analysis. This suggests that the mutational status of PTC is associated with fibrosis, but also the density of CAFs and their activation.

We showed that PTC^V600E^ have a higher expression of proliferation-associated mRNA and of the Ki67 protein, and lower expression of thyroid differentiation genes. Potential links have been described between aggressiveness and the tumor-stroma cross-talk in PTC. One group proposed that PTC^V600E^ could be more invasive, *via* a decrease of CDH1 expression [28] and another suggested that PTC^V600E^ were more susceptible to TGF-β-induced EMT through a MAPK-dependent process [29]. However, stromal fibroblasts and endothelial cells could also express EMT markers [30]. And we showed here the high fibroblast content of PTC^V600E^. Hence, results relying on expression of EMT-markers should be interpreted with caution.

Altogether, our results depict a more aggressive phenotype of PTC^V600E^, helped by denser, more active and proliferative CAF population, i.e. a higher desmoplastic reaction associated with BRAF^V600E^ in PTCs. These tumors could in theory be treated accordingly. Approved drugs targeting the BRAF-mutated pathway would represent good candidates but, although at an early stage of clinical assessment, thyroid cancers might respond to a lesser degree to BRAF-inhibitor compared to melanoma and other cancer types presenting the mutation [31]. Interestingly, higher CAFs content and stiffer extracellular matrix (ECM) have been shown to protect melanoma^V600E^ cells against BRAF-pathway inhibitors [32]. The higher density and activation of CAFs in PTC uncovered here could be responsible for a resistance, as exampled in melanoma [32]. Hence, our results would be compatible with improved treatment response via combination of ECM and BRAF-inhibitors in PTC^V600E^ [32].

## Materials and Methods

**PTC^V600E^ case description.** A 51-year-old female was diagnosed with a bilateral PTC (right lobe: 0,6 cm; left lobe: 2,8cm) with concurrent metastatic involvement of the lymph nodes (stage IVA, pT3N1b). She was treated with a total thyroidectomy and a lymphadenectomy, with 5 nodal metastases in the recurrent and mediastinal areas (Figure 2A). Iodine-131 radiotherapy was administered one month after surgery. Twelve months after thyroidectomy, she was operated for nodal recurrences (4 nodal metastases from the left cervical, 3 of which were included in the study, annotated M2, M3 and M4). Histopathological examination revealed a classical PTC developing in a goitre context, and suggested a tumour cell fraction greater than 70%.

**Tissue samples.** For immunohistochemistry (IHC) and exomes analyses, tumour and normal thyroid tissues were obtained from Jules Bordet Institute tissue bank (Brussels, Belgium). We have studied 4 foci of the primary PTC from the left lobe (T1-T4), 1 from the right lobe (T5) and 4 nodal metastases (M1-M4) (Figure 2A). Non-cancerous adjacent tissues from the tumour (N1-N3), the lymph node (LN), and a blood sample were taken as controls. Immediately after surgery, tissues were macro-dissected in several blocks. Each block was cut into two equal parts. One part was embedded in paraffin for histological analyses and the other part immediately snap-frozen in liquid nitrogen and stored at -80°C until RNA and DNA extraction processing (Figure 2B). Only DNA was analysed for this study.

**Extraction, quality assessment of nucleic acids and reverse transcription.** DNA from tumour and normal thyroid tissues was extracted using the DNeasy Blood and Tissue Kit (QIAGEN, Hilden, Germany) according to the manufacturer’s recommendations. DNA concentrations were spectrophotometrically quantified, and nucleic acids integrity was checked using an automated gel electrophoresis system (Experion, Bio-Rad, Hercules, USA).

**Sanger sequencing of DNA and cDNA samples.** Primer sequence for BRAF DNA FWD: TGCTTGCTCTGATAGGAAAATG; REV: CCACAAAATGGATCCAGACA. To look for mutations, PCR reactions were performed on 80 ng of amplified genomic DNA or 2 μl of cDNA using the recombinant Taq DNA polymerase kit (Invitrogen, Carlsbad, California, United States) and appropriate primer pairs (Supplementary material). PCR products were purified with the QIAquick PCR Purification Kit (QIAGEN, Hilden, Germany) according to the manufacturer’s instructions. Sequencing was performed with the BigDye Terminator V3.1 Cycle Sequencing Kit (Applied Biosystems, Foster City, USA) on the sequencer ABI PRISM 3130 (Applied Biosystems, Foster City, USA) using the genetic analysis program 3130-XI.

**Exome sequencing.** Exome DNA-sequencing was performed with the Nextera Rapid Capture Expanded exome kit (Illumina Inc., San Diego, California, USA). All material was sequenced with the Illumina HiSeq 2000 platform (Illumina Inc., San Diego, California, USA) using 100 base-pair paired-end reads for a yield of ∼180 million reads per sample.

**Sequence analyses.** Duplicates were removed using Picard’s MarkDuplicates (http://picard.sourceforge.net) with default parameters. Reads were mapped to hg19 using BWA [33] with default settings. Reads were realigned locally with the GATK IndelRealigner (v1.4-15) [34]. Somatic mutations were called with MuTect [35] in both tumour and normal adjacent tissues, with default parameters, taking the blood tissue as normal reference (Supplementary material).

**Mutation calling and genotyping.** All genomic positions returned by MuTect were pooled in a single variant call format file. For each position, we retrieved the coverage and allelic ratio of each position in all BAM files. Mutations detected in at least one tumour sample were further filtered based on the coverage and the allelic ratios in the pool of non-cancerous tissues (blood, non-cancerous thyroid N1-N3 and nodal tissue LN) and the pool of tumour tissues (T1-T5 and M1-M4). Minimal total coverage of 100 aligned reads and 0 mutated reads in normal pool; minimal total coverage of 100 aligned reads and at least 1 mutated read in the tumour pool. The latter is a conservative threshold, well suited for the subsequent analysis (see Figure 2G).

**NGS and Sanger allelic fractions.** BRAF^V600E^ DNA allelic fractions were measured using 2 independent techniques: Sanger sequencing and Illumina HiSeq2000 sequencing.

For Sanger sequencing, *ab1* files were loaded in *R v3.1.0* with the package *seqinr v3.0-7* [36]. First we corrected for the background offset, i.e. subtracting the average of the 4 channel median values from the signals. The 2 peaks of interest were manually identified at the V600E position c.1799T>A. Fractions were then assessed dividing the height of the peak corresponding to the mutation by the total height of the two peaks.

Illumina sequencing fractions were defined as the ratio of mutated aligned reads and the total aligned reads retrieved from the BAM files. Allelic fractions for all other somatic mutations were compared to allelic ratio for BRAF^V600E^ in each sample. For each sample only allelic fractions greater than zero were considered.

**IHC ratio.** For each block, one slice was stained with the antibody specific for the BRAF^V600E^ protein. Scanned images were saved as *png* files of lower resolution with *NDP.view2* (www.hamamatsu.com/jp/en/U12388-01.html). Pixels were set to white if they passed manually set thresholds on colour hue, saturation and value (H, S, V). Colour hues of the stained and unstained tissues were around 260 and 240, respectively. Hence the colour hue threshold was set to 250. The other pixels were set to black. Two binary images were derived: *image1* with the total tissue in white (0<=H<=360; S>0.1; V>0.2) and *image2* with the stained part of the tissue in white (250<H<=360; S>0.1; V>0.2). The binary images were eroded to remove noise from the background: isolated pixels, i.e. pixels with less than 60% marked pixels in a 5-pixel radius, were set to black. Finally, we define an IHC ratio from image analysis, corresponding to the ratio of mutated-protein expressing tissues on the total bulk tissue, i.e. the IHC ratio is the ratio of number of white pixels in image2 and image1.

**Immunohistochemistry.** IHC staining was performed on 8 μm thick sections prepared from paraffin-embedded tissues with the primary antibody directed against BRAF^V600E^ (Spring Bioscience, Pleasanton, USA; Clone VE1) or Ki67. The sections were deparaffinised, pre-treated with CC1 (EDTA, pH8.4) and incubated with the antibody at a 1:100 dilution. The revelation was performed with detection kit ref. 760-501 (Roche, Vilvoorde, Belgium).

**3D reconstitution.** For two of the paraffin tumour blocks, T1 and M1, over 100 adjacent slices were cut and stained for BRAF^V600E^ protein. These images were used to reconstruct the associated 3D volumes of the stained parts, i.e. the tumour (Figure 2F shows that mutated tumour cells express the mutated protein; Figure 2G shows that all tumour cells are mutated). 3D reconstitution was obtained using *BioVis3D* (biovis3d.com). Images of adjacent slices were manually aligned. Then, on each slice, contours of the stained tumour parts were manually created and were either linked, divided or merged from slice to slice. Finally BioVis3D rendered a 3D model of these connected 2D contours (Supplementary Video S1).

**Ki67 quantification.** For each block, proliferation rates were calculated on scanned images stained for Ki67 by manually counting the Ki67 stained nuclei against the non-stained nuclei, which is a reliable method to derive Ki67 index [37]. Ki67 indices were computed separately in multiple regions for the stromal fibroblasts and follicular parts, not necessarily hot spots of high proliferation. This approach provided average proliferation rates and associated estimates of the measurement error across the slice.

**Independent PTC set.** Twenty-two of the most recently added PTC from Jules Bordet Institute (Brussels, Belgium) were taken as independent set for H&E, Ki67 and BRAF staining. They were selected on the basis of sample age and thus constituted a mixture of PTC subsets, BRAF mutant (N=13) and wild type (N=9). **TCGA data.** We retrieved PTC RNAseq2 expression data from TCGA. Mutation annotated files v0.2.0 were downloaded from the Broad Institute Firehose website (https://confluence.broadinstitute.org/display/GDAC/MAF+Dashboard). Mutation profiles were available for 366 PTC, from which 211 harbored BRAF^V600E^. Histological images stained for haemotoxylin and eosin (H&E) corresponding to TCGA unique ID were manually downloaded from the cancer image archive for thyroid, colorectal and skin cancers (http://cancer.digitalslidearchive.net/).

**PCA and gene signatures.** Principal component analysis (PCA) projects all transcriptomic profiles on the two axes (first components) that maximise total observed variance in the data. This allows intuitive visualisation of complex multidimensional data. To better interpret the clustering of the samples in the PCA space, five gene signatures were projected together with the samples: *Thyroid, Stroma, T-cell, Proliferation* and *BRAF-RAS,* indicating the direction of their higher expression. Axes of gene signatures were computed as projections of sample vectors with 1-values for genes belonging to the signature, 0-values otherwise. They are depicted as arrows from the centre of the data to the projected position, as computed by principal component analysis (PCA). The lengths of the arrows were normalised to 1, and then multiplied by the weighted sum of two Spearman correlation coefficients: across all samples, we compute the two correlations between the first or second PCA component coordinates and the medians of expression of the genes from the signature. They are weighted by the percentage of variance explained by each axis. The thyroid signature was made of 16 thyroid-specific genes described in the TCGA publication [15]. The signatures *Stroma* and *T-cell* were each composed of the 50 genes that were most correlated with *PLAU* and *TRAC* genes, respectively, as computed by Gemma [38] in a master human dataset of 2160 microarray profiling studies. The *BRAF-RAS* signature was made of the 71 most differentially expressed genes between PTC^V600E^ and PTC harbouring mutation in the *RAS* family of genes in TCGA, similarly to the signature developed in the TCGA publication [15]. Differential expression was assessed with the Rank Product methodology [39] as implemented in RPlite (sourceforge.net/projects/rplite/). The 71 genes with best scores were kept in the signature. The proliferation signature was made of the 101 genes whose expression most correlated with *PCNA* expression, i.e. a marker of proliferation, in a microarray dataset of normal human tissues, as described earlier [40].

**Data availability.** Sequence data are available in The European Genome-phenome Archive under accession number EGAB00000001141. All scripts and additional data are available upon request.

**Study approval.** The patient gave written informed consent. The medical ethics committee of the Jules Bordet Institute gave approval for this study (The Medical Ethics Committee of Institut Jules Bordet – LEC; 07/06/2012; ref. as-1978).

## Author contribution

AA processed the PTC sample for next-generation sequencing. AA performed Sanger sequencing experiments. MT and DG conducted bioinformatic analyses. MT performed image analyses. AA and MT reconstructed the 3D microvolumes. MT, AA, VD, CM interpreted the results. MT, VD, CM, and JED wrote the manuscript. VD, CM, and SLP designed the experiments. GA collected patient samples. DL, LC and NDSA collected and reviewed the histopathological slices of the independent PTC dataset and the sequenced PTC case. AA, IL and TC performed immune stainings.

## Acknowledgments

Some of the results presented here are based upon data generated by the TCGA Research Network: http://cancergenome.nih.gov/. The authors wish to thank Chantale Degraef for her excellent technical assistance. MT was supported by a FRIA/FNRS grant. This work was supported by the Fonds de la Recherche Scientifique FNRS-FRSM, WELBIO, Plan Cancer Belgique, les Amis de l’Institut Bordet, and Fondation Van Buuren. VD was funded by FNRS, grant J009714F.

## Notes

**Conflict of interest** The authors declare no conflict of interest. This original work has not been submitted or published elsewhere

